# Improved genome assembly of double haploid *Prunus persica* siblings ‘Lovell 2D’ and ‘Lovell 5D’ and the peach NLRome

**DOI:** 10.1101/2025.06.03.657495

**Authors:** Christopher Gottschalk, Jordan R. Brock, Ben N. Mansfeld, Dorrie Main, Sook Jung, Ping Zheng, Cheryl Vann, Mark Demuth, Dennis Bennett, Zongrang Liu, Chris Dardick

**Affiliations:** USDA-ARS Appalachian Fruit Research Station, 2217 Wiltshire Rd., Kearneysville, WV, 25430; Washington University in St. Louis, 1 Brookings Dr., St. Louis, Missouri, 63130; Washington State University, 255 E Main St, Pullman, WA 99163

## Abstract

*Prunus persica* (peach) has long served as a model fruit tree for studying phenological events. It has a relatively small genome and exhibits tremendous plasticity in climate tolerances due to the high variation of chill requirements, bloom times, and fruit ripening times. The peach variety ‘Lovell 2D’ was used to generate one of the first high-quality genome assemblies for a tree species, using Sanger sequencing of genetic-map ordered BAC clones. A key to the high quality of this early assembly was the use of a doubled haploid variety, which eliminates the challenges posed by mixed haplotypes. Here, we re-sequenced and assembled the ‘Lovell 2D’ genome along with a doubled haploid sibling ‘Lovell 5D’ using 3rd generation technologies. The resulting genomes were significantly more contiguous than the current ‘Lovell 2D’ reference genome (ver2.0 updated in 2017) and are closer to the estimated total genome size for peach (265Mb). In addition, new gene, transposable element (TE), and Nucleotide-binding domain and Leucine-rich repeat receptor (NLR) annotations were performed to enhance the integrity and utility of the genome. These updated peach doubled-haploid reference assemblies will provide the research community with an improved reference genome for genomics-guided studies and breeding efforts.

## Introduction

Peach (*Prunus persica* L. Batsch) is a temperate tree fruit crop adapted to diverse climates across the globe. U.S. production of peach ranges between 650,000 - 710,000 tons per year and routinely experiences crop losses due to extreme weather events (i.e., spring frost events) (Foreign Agricultural Service USDA, 2023). Moreover, production of peach is threatened by numerous diseases such as Brown Rot (caused by *Monilinia spp*.), which are becoming more prevalent in response to the climacteric conditions (Luo *et al*., 2022). Improved genomics tools can aid in the development of more secure, sustainable, and resilient peach varieties. The first peach genome released was of ‘Lovell 2D’, a doubled haploid variety that was assembled from first-generation paired-end Sanger sequencing of fosmid and BAC clones (Verde *et al*., 2013). The resulting assembly had 234 scaffolds placed into eight pseudomolecules using a genetic map. The final assembly was 224.6 Mb in length with a scaffold N50 of 4 Mb (Verde *et al*., 2013). A revision was performed four years later using Illumina next-generation sequencing data (Verde *et al*., 2017). This revision effort resulted in the closing of 212 gaps in the ‘Lovell 2D’ assembly with a minor sequence gain of ∼25 Kb (Verde *et al*., 2017). 2021 saw the release of four new peach genomes based on PacBio third-generation sequencing platform. The first was for the Chinese variety ‘Longhua Shui Mi’ with a reported Contig N50 of 5.17 Mb and a length of 257.2 Mb (Yu *et al*., 2021). The second genome was from a Chinese flat peach variety ‘124 Pan’ (Zhang *et al*., 2021).

This assembly had a reported N50 of >26 Mb and a total length of 206 Mb. The third genome was for another Chinese cling-stone variety with a reported scaffold N50 of 29.68 Mb and a length of 247.33 Mb (Cao *et al*., 2021). That assembly represented 99.8% of the estimated genome size, the closest to date. The fourth genome was for a semi-dwarf breeding line variety ‘Zhongyoutao 14’ and the assembly had a reported scaffold N50 of 27.89 and a total length of 228.82 Mb (Lian *et al*., 2022).

Over the past decade, the peach ‘Lovell 2D’ reference genome has served as a critical resource to the research community. Although tremendous efforts have been made to improve the genome resources for peach, little has been done since 2017 to improve the reference genotype’s assembly. To address this, we sequenced and assembled an improved peach reference genome using the original double haploid ‘Lovell 2D’ and a sibling doubled haploid ‘Lovell 5D’. To achieve high-quality genome assemblies, we utilized the cutting-edge Oxford Nanopore PromethION sequencing platform to generate high accuracy simplex reads and combined them with PacBio HiFi reads. Combining these sequencing data types with the latest genome assembly software that utilizes mixed data types resulted in near-complete genome assemblies. These updated doubled haploid reference genomes will provide a valuable resource for molecular breeding strategies helping overcome apparent domestication bottlenecks that may limit breeding germplasm diversity for certain critical traits (Verde et al., 2013).

## Materials and Methods

### Plant Materials

Field grown trees of ‘Lovell 2D’ and ‘Lovell 5D’, of an estimated ∼30 years of age, are maintained in the breeding germplasm at the Appalachian Fruit Research Station (AFRS) in Kearneysville, WV (Fig. 1A). The trees were grown using standard commercial practices for training, disease, and pest management. In 2017, young leaf material was collected and flash frozen. The tissues remained frozen at -80°C until removed for DNA extraction. ‘Lovell 2D’ and ‘Lovell 5D’ are available through the National Plant Germplasm System as DPRU 263 – ‘LOV-2-Haploid’ and PI 673461 – ‘LOV-5-Haploid’, respectively (Fig. 1B,C).

**Figure 1.**
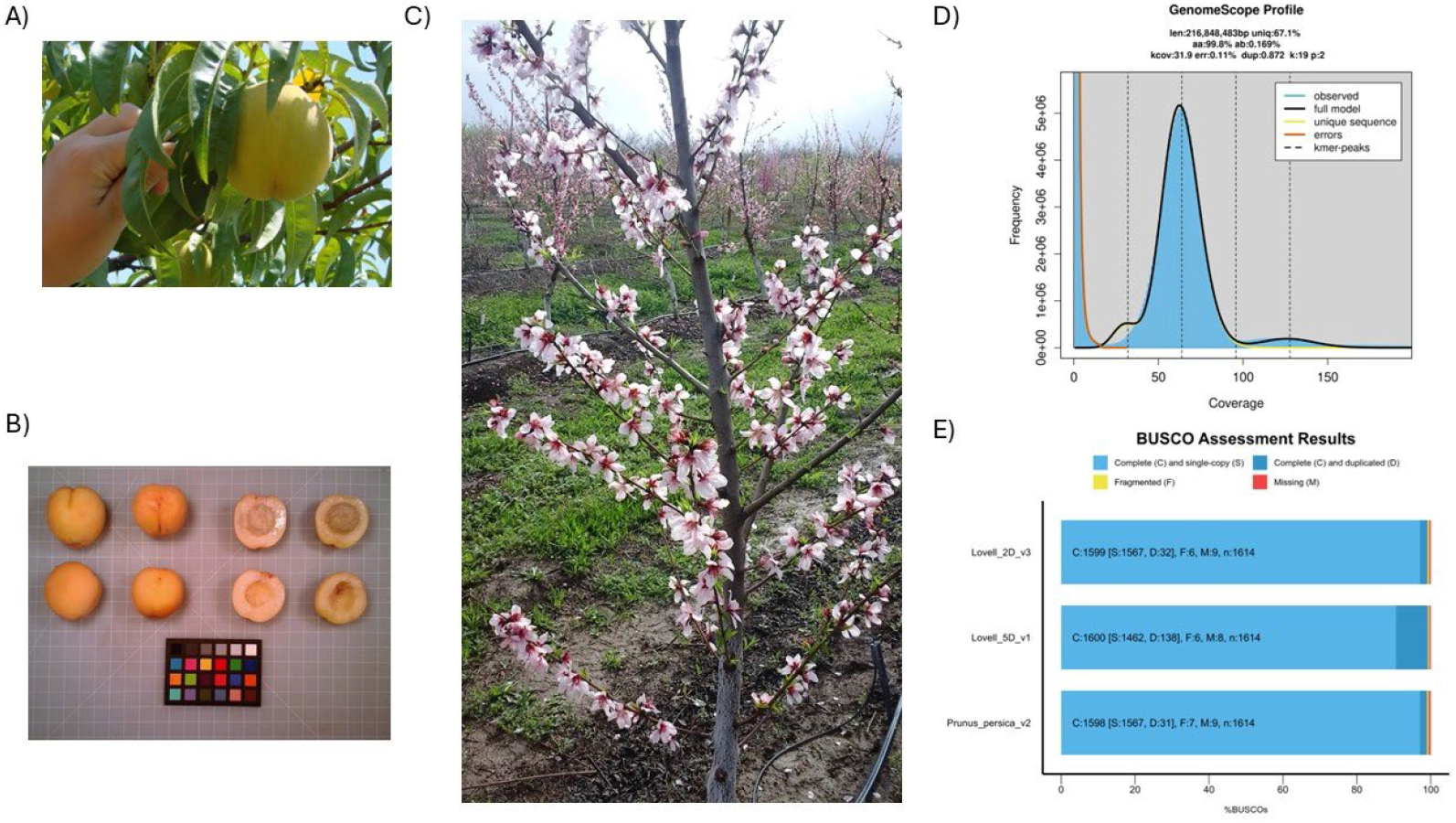
A) ‘Lovell 5D’ fruit grown at AFRS. B) ‘Lovell 5D’ (PI 673461) fruit collected from the USDA germplasm in Parlier, CA. C) ‘Lovell 5D’ (PI 673461) tree in bloom at the USDA germplasm in Parlier, CA. D) GenomeScope 2.0 *k-*mer frequency plot and estimated genome statistics for ‘Lovell 5D’. E) BUSCO genome plots for the ‘Lovell 2D v3.0’, ‘Lovell 5D v1.0’, and ‘Lovell 2D’ v.2.0.a1 reference genome. Photos of ‘Lovell 5D’ (PI 673461) were retrieved from the USDA National Plant Germplasm System.

### DNA Extraction

Frozen leaf tissues were ground into a fine powder using a mortar and pestle chilled with liquid N_2_. A total of ∼1 gram of tissue was used per extraction. High-molecular weight (HMW) DNA was extracted using the LeafGo (leaf to genome) extraction protocol that relies on gravity-based filter columns (Qiagen Genomics Tip Kit, Germantown, MD) (Driguez *et al*., 2021). There were no modifications to the protocol except during the precipitation of HMW DNA using 2-propanol. Following the addition of 2-propanol, the HMW DNA sample was centrifuged at 5,000g for 15 minutes with the centrifuge chilled to 4°C. Following pelleting, the HMW DNA pellet was washed using 80% ethanol and centrifuged at 5,000g for 15 minutes with the centrifuge chilled to 4°C to pellet the DNA. This washing step was repeated twice and after the second wash the pellets were left to air dry for 15 minutes. Following drying, HMW DNA was dissolved in TE buffer. The resulting HMW DNA samples were tested for fragment size, quality, and purity using an Agilent Tapestation (Santa Clara, CA) using the Genomic DNA screen tape and reagents according to manufacturer protocols.

### Library preparation and sequencing

Oxford Nanopore Technologies (ONT, Oxford, UK) sequencing libraries were prepared for each variety using the Ligation Sequencing Kit V14 (SQK-LSK114) without modifications. The library DNA quality and quantity were measured using an Agilent Tapestation with the Genomic DNA screen tape and reagents. Libraries were sequenced on the ONT PromethION platform using independent R10.4.1 flow cells. Basecalling was conducted within the MinKnow GUI software (v. 23.11.7) using the Dorado basecaller (v.7.2.13) for real-time simplex basecalling. Following the completion of sequencing, command-line Dorado (v.0.7.2) was executed a second time with the dna_r10.4.1_e8.2_400bps_sup@v5.0.0 model to generate highly accurate >Q20 reads. The resulting simplex reads were evaluated for PHRED scores and length using Nanoplot (v.1.42.0; De Coster and Rademakers, 2023). The reads were then filtered for a length of 40 Kb for ‘Lovell 2D’, and a length of >20 Kb and Q25 for ‘Lovell 5D’ using Filtlong (v.0.2.1; https://github.com/rrwick/Filtlong). For ‘Lovell 2D’, we additionally executed the command-line Dorado (v.0.4.1) a second time using the ‘duplex’ parameter with the dna_r10.4.1_e8.2_400bps_hac@v4.2.0 model to generate duplex reads. For ‘Lovell 5D’, we used the same HMW DNA sample to sequence on a PacBio Revio (Menlo Park, CA) Flow Cell to generate a HiFi read dataset.

### Genome assembly, scaffolding, and annotation

To estimate genome size and to confirm the doubled haploid state of ‘Lovell 5D’, we counted the *k-mers* within the HiFi dataset. The *k-*mer counting was performed using KMC (v.3.2.4) using the following parameters: -k19 -t20 -m60 -ci1 -cs10000 as recommended by GenomeScope 2.0 (http://genomescope.org/genomescope2.0/) (Kokot *et al*., 2017; Ranallo-Benavidez *et al*., 2020). The counted *k-*mers were then transformed into a histogram using the kmc_tools script in KMC package with the transform reads histogram reads.histo -cx10000 parameters (Kokot *et al*., 2017; Ranallo-Benavidez *et al*., 2020). The resulting histogram was then uploaded to GenomeScope 2.0 for *k-*mer plot generation (http://genomescope.org/genomescope2.0/) (Ranallo-Benavidez *et al*., 2020).

We assembled the genome of ‘Lovell 2D’ using two independent assembly programs. First, we used Verkko (v.2.0) with the duplex nanopore data used as HiFi input and the simplex data as ultra-long reads (Rautiainen *et al*., 2023). All other parameters were set to default. Our second approach utilized HiFiasm (v.0.19.8) with the duplex data used as HiFi input and the simplex data as ultra-long reads (Cheng *et al*., 2022). Again, all other parameters were set to default. The resulting assemblies were analyzed for completeness using GenomeTools statseq (v.1.6.5) (Gremme *et al*., 2013). Based on the statistics of each assembly, the most contiguous and longest ‘Lovell 2D’ assembly was selected to move forward with. Our assembly approach for ‘Lovell 5D’, used HiFiasm (v.0.19.9) with the HiFi sequence data used as HiFi input and the Oxford Nanopore simplex data as ultra-long reads (Cheng *et al*., 2022). We invoked the parameters --ul-cut 20000, --telo-m TTTAGG and -l0 with HiFiasm. The resulting assemblies were analyzed for completeness using GenomeTools seqstat (v.1.6.5) (Gremme *et al*., 2013). The resulting primary assembly fastas for each genotype were scaffolded using RagTag (v2.1.0; Alonge *et al*., 2022).

This scaffolding method relied on mapping the genomes of each genotype to a reference, and in our case we used the ‘Lovell 2D’ v2.01.a1 scaffolded reference assembly (Alonge *et al*., 2022; Verde *et al*., 2017). The agp file generated by RagTag was then converted into a fasta using agptools assemble (https://github.com/WarrenLab/agptools). Final scaffolding stats were then obtained using gfastats (v.1.3.7) (Formenti *et al*., 2022).

The scaffolded assemblies were then annotated for transposable elements (TE) and repeats using EDTA (v.2.0.1) (Ou *et al*., 2019). Here, we used a highly curated TE library from the ‘Texas’ almond genome and CDS sequences from the ‘Lovell 2D’ v.2.0.a1 genome as input into EDTA for *de novo* TE/repeat annotation (Castanera *et al*., 2024; Verde *et al*., 2017). The resulting annotation and genome assembly were evaluated for quality using Long-Terminal Repeat (LTR) Assembly Index (LAI) within the EDTA suite (Ou *et al*., 2018). EDTA also produced a masked genome file, which we used as input into the gene annotation pipeline.

For gene annotation, we utilized the MAKER pipeline, which incorporates using transcriptional and protein homology evidence (Holt and Yandell, 2011). For transcriptional evidence, the primary transcripts from ‘Lovell 2D’ v2.0.a1 genome were downloaded from Genome Database for Rosaceae (GDR) (Verde *et al*., 2017; Jung *et al*., 2019). For protein evidence based on homology, we downloaded and used the Araport11 protein sequence dataset (Cheng *et al*., 2017). We performed three rounds of MAKER (v.3.01.03) annotation as previously performed by the authors (Mansfeld *et al*., 2023). In short, the first round was the evidence-based round with *ab initio* gene prediction training. The following two rounds of annotation were conducted using retrained *ab initio* gene prediction and carrying over of evidence-based annotations. The gene predictors used were SNAP (v.2013_11_29) and Augustus (v.3.5.0) (Korf, 2004; Hoff and Stanke, 2019). Noncoding genes were predicted using Infernal (v.1.1) (Nawrocki and Eddy, 2013). Genome and transcriptome quality were additionally evaluated using BUSCO (v.5.8.0) with the embryophyta odb10 (Manni et al., 2021).

### Annotation of R-genes

We set out to annotate the *R-*gene space in our two new genome assemblies along with the four previously released *P. persica* genomes hosted on the GDR webpage (Verde et al., 2017; Jung *et al*., 2019; Cao et al., 2021; Zhang et al., 2021; Lian et al., 2022). These additional genomes included: ‘Lovell 2D’ v2.0.a1 (Prup; Verde et al., 2017), ‘Chinese Cling’ (Ppcc; Cao et al., 2021), ‘Zhongyoutao 14’ (CN14; Lian et al., 2022), ‘124 Pan’ (124Pan; Zhang et al., 2021). Each genome had its representative sequence processed to remove any soft masking. We then ran each genome separately through the FindPlantNLR snakemake pipeline (v.2.0) following the software’s default methodology (Chen et al., 2023). The resulting annotations of the complete catalog of *R-*genes (NBARC) for each genome was then compared to its respective gene annotation using gffcompare (v.0.12.6) to identify the number of novel annotations using the “u” flag (Pertea and Pertea, 2020).

### Comparative genomics

Comparative genomics was conducted using the SyRI (v.1.7.0) package (Goel et al., 2019). Default parameters were used when executing the SyRI commands as described in the manual. The plotting of the syntenic regions was generated using the plotsr (v.1.1.0) program as recommended in the SyRI manual (Goel et al., 2019).

### Pan-NLR analyses

Nucleotide-binding domain and Leucine-rich repeat receptors (NLR) gene synteny of the primary (or longest) isoforms of NLRs were determined between the six accessions of *Prunus persica*, using the genome of ‘Lovell 2D v3.0’ as a reference genome for comparison. Pairwise analyses of syntenic genes and NLRs were conducted between all genomes. Coding sequences were extracted from genome annotations using gffread v0.12.7 (Pertea and Pertea, 2020). Macrosynteny and NLR gene synteny were assessed using the Python version of the MCScanX pipeline (v1.1.12) (Tang et al., 2008; Wang et al., 2012). Macrosynteny was defined with a minimum syntenic block size (−-minspan) of 10 genes for both sets of analyses. Syntenic blocks and collinear gene pairs were identified, and results were visualized using the jcvi.graphics.karyotype module.

## Results and Discussion

We generated a total of 69.44 Gb of ONT simplex sequence data for ‘Lovell 2D’. This sequencing yield represents an estimated 262x coverage, assuming the estimated genome size is 265 Mb (Verde *et al*., 2013). Following length and quality filtering, the ‘Lovell 2D’ simplex data was reduced to 5.7 Gb with a N50 of 47.6 Kb and a median read quality of Q17.4. The ONT sequencing of ‘Lovell 2D’ additionally yielded 5.53 Gb of duplex reads that represent 21x coverage with a median read quality of Q24.8. We generated a total of 86.63 Gb of ONT simplex sequence data for ‘Lovell 5D’. This sequencing yield represents an estimated 327x coverage. Following length and quality filtering, the ‘Lovell 5D’ simplex data was reduced to 24.02 Gb with a N50 of 20.7 Kb and a median read quality of Q26. The HiFi sequencing of ‘Lovell 5D’ yielded 10.51 Gb of HiFi reads with a 6.33 Kb mean length and a median read quality of Q39. This amount of sequence data was deemed sufficient to proceed with genome assembly for both genotypes. We counted *k-*mers in the HiFi data to confirm the double haploid state of ‘Lovell 5D’. GenomeScope 2.0 reported a homozygosity minimum rate of 99.81% and an estimated genome size of 216 Mb (Fig 1D). This lower estimated genome size than the previous estimate of 265 Mb could be an artifact of the homozygosity of ‘Lovell 5D’ and the relatively short PacBio read lengths (Verde *et al*., 2013).

### An improved ‘Lovell 2D’ genome

Verkko’s initial assembly of ‘Lovell 2D’ yielded a primary assembly of 267.6 Mb contained in 969 contigs (Table 1). The contig N50 was 16.5 Mb, which is 64-fold improvement in contig length and ∼4-fold improvement in scaffold lengths over the ‘Lovell 2D’ v.2.0.a1 reference genome (Verde *et al*., 2017).

Using the ‘Lovell 2D’ v.2.0.a1 genome for homologous scaffolding with RagTag generated 904 scaffolds with a scaffold N50 of 28.2 Mb. The scaffold N50 is nearly 1 Mb larger than the ‘Lovell 2D’ v.2.0.a1 reference genome (Verde *et al*., 2017). We also observed a small increase in complete BUSCO percentage of 0.1% over the ‘Lovell 2D’ v.2.0.a1 reference genome (Verde *et al*., 2017). However, we observed an increase of 27.7% to the LAI score compared to the ‘Lovell 2D’ v.2.0.a1 reference genome (Verde *et al*., 2017) (Table 1). Increased LAI indicates more contiguous assembly through the complex, repetitive regions of the genome.

### A new genome for ‘Lovell 5D’

HiFiasm generated a primary assembly of 257.8 Mb contained in 400 contigs (Table 1). The contig N50 was 23 Mb, which is 90-fold increase in contig length and >5-fold increase in scaffold length over the ‘Lovell 2D’ v.2.0.a1 reference genome (Verde *et al*., 2017). Using the ‘Lovell 2D’ v.2.0.a1 genome for homologous scaffolding with RagTag generated 193 scaffolds with a N50 of 29 Mb. The scaffold N50 is nearly 2 Mb larger than the ‘Lovell 2D’ v.2.0.a1 reference genome (Table 1) (Verde *et al*., 2017). Our assembly is the closest reported to the predicted length from Verde *et al*. (2013) of 265 Mb and exceeded the estimated length from our *k-*mer analysis (Fig. 1D). We observed no difference in complete BUSCO percentage over the ‘Lovell 2D’ v.2.0.a1 reference genome (Verde *et al*., 2017).

However, we observed a nearly 7% increase in the complete duplication rate of BUSCOs. Lastly, we observed an increase of 23.9% in the LAI score compared to the ‘Lovell 2D’ v.2.0.a1 reference genome (Verde *et al*., 2017) (Table 1).

**Table 1. Genome assembly statistics for the ‘Lovell 2D’ v3.0, ‘Lovell 5D’, and the reference ‘Lovell 2D’ genome**.

### Comparison of gene annotations

The annotation of transposable elements and other repetitive features for our two genomes was found to be lower than the reported amount in the ‘Lovell 2D’ v.2.0.a1 reference genome (Verde *et al*., 2017) (Table 2). Both of our genomes were found to have 82.9 – 85.1 Mb of repeat sequences compared to 84.4 Mb in the ‘Lovell 2D’ v.2.0.a1 reference genome. This lower content represents a decrease of ∼4-7% over ‘Lovell 2D’ v.2.0.a1. However, the higher LAI scores for our assemblies indicate improved contiguity and assembly quality of these repetitive regions. We observed ‘Lovell 5D’ exhibiting higher repeat space. This increase in repeat space was observed within all the annotated features except unknown LINE elements (Table 2).

We undertook a *de novo* gene annotation approach as opposed to lifting over the annotation from ‘Lovell 2D’ v.2.0.a1. We identified 26,834 and 26,764 genes in the new ‘Lovell 2D’ and ‘Lovell 5D’ genomes, respectively (Table 2). These results are lower than ‘Lovell 2D’ v.2.0.a1 by 37 and 99 genes, respectively. The resulting complete BUSCO scores reflected this, with the new ‘Lovell 2D’ being 97.00% and ‘Lovell 5D’ being 98.40%. Whereas ‘Lovell 2D’ v.2.0.a1 exhibited a 99.10% complete BUSCO score. Although lower by <100 genes from the reference, these *de novo* gene annotations were conducted with no manual curation, unlike the reference. We also conducted a more modern approach to annotation for ncRNAs in our genomes, which resulted in a more thorough annotation compared to ‘Lovell 2D’ v.2.0.a1. These two new resources now include annotated ncRNA features such as rRNA, miRNA, and snoRNA (Table 2).

**Table 2. Genome annotation statistics for the ‘Lovell 2D’ v3.0, ‘Lovell 5D’, and the reference ‘Lovell 2D’ genome**.

### Synteny among ‘Lovell’ genomes

We conducted a syntenic analysis between our two new genomes and the ‘Lovell 2D’ v.2.0.a1 reference genome (Verde *et al*., 2017) (Fig. 2). Overall, synteny was highly conserved with >96% of the reference genome being found in 593 syntenic regions with the v3.0 assembly of ‘Lovell 2D’ (Table 3). Within this comparison, we also identified >7,000 insertion events corresponding to ∼480 Kb of sequence, which is less than 2% of the total newly added sequence. We also identified several structural variations between the two assemblies. Of which, 10 were inversions, 65 were translocations, and 1,256 were annotated as duplications. The ‘Lovell 5D’ assembly also exhibited a high level of conserved sequence identity and organization with its sibling, the ‘Lovell 2D’ reference assembly and our third version (Fig. 2). Comparing the ‘Lovell 2D’ v3.0 with ‘Lovell 5D’, we identified 70 syntenic regions that span 98.8% of the total chromosomal length of ‘Lovell 2D’. We also observed lower structural variation (inversions and translocations) and sequence annotated variation in this comparison than within the intra-synteny of the ‘Lovell 2D’ assemblies. However, the total duplicated sequence was 1.5Mb more than the intra-synteny of the ‘Lovell 2D’ assemblies (Table 3; Fig. 2). This result was also observed in the higher rate of complete and duplicated BUSCO genes in the ‘Lovell 5D’ assembly. These results could be attributed to the higher quality of data (PacBio HiFi + ONT long reads) used in the assembly ‘Lovell 5D’, which resulted in it exhibiting greater contiguity and BUSCO scores (Table 1).

**Figure 2.**
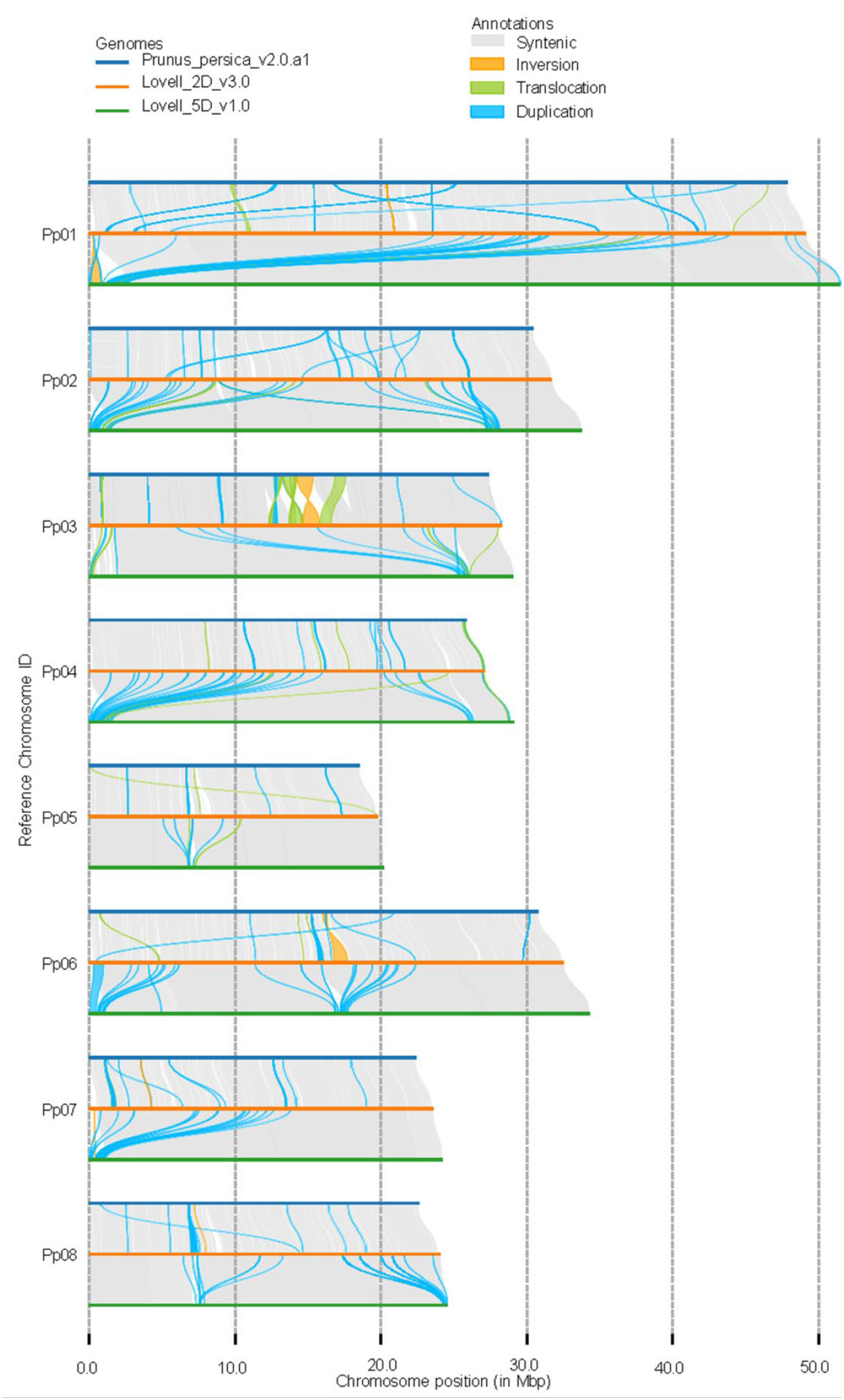
Structural variation and synteny between ‘Lovell 2D’ v3.0, ‘Lovell SD’, and the reference ‘Lovell 2D’ genome.

**Table 3. Genomic variation statistics for the ‘Lovell 2D’ v3.0, ‘Lovell 5D’, and the reference ‘Lovell 2D’ genome**.

### R-gene space within P. persica genomes

Like other crops, peach production can suffer from several plant pathogens, including but not limited to peach leaf curl, brown rot, and peach scab (Luo *et al*., 2022). Understanding the repertoire and diversity of pathogen resistance loci in the currently sequenced peach genomes is crucial for mobilizing these genes in future germplasm improvement. Using the FindPlantNLR pipeline, we identified between 343 - 448 Nucleotide Binding Leucine Rich repeat domain (NLRs or *R*-genes)*-*genes (primary transcripts) in the *P. persica* genomes (Chen et al., 2023). The Chinese Cling genome exhibited the highest *R-*gene count with 448, of which 131 were not overlapping with the reference gene annotation (Cao et al. 2021) (Table 4). The ‘Lovell’ double haploid clones 2D and 5D contained 382 and 401 *R*-genes, respectively, for the latest versions. The number of novel *R-*gene loci in the *R-*gene annotation was between 43 - 53 for the ‘Lovell’ genomes. In comparison, the ‘Lovell 2D’ v.2.0.a1 genome had 471 *R-*genes annotated with the FindPlantNLR pipeline, and only 28 were novel compared to the reference gene annotation (Table 4) (Verde et al., 2017).

Syntenic analysis of NLR genes revealed that NBARC-containing loci are largely colinear with broader genome-scale macrosynteny, indicating minimal NLR-specific genome rearrangement across accessions (Figure 3A). However, some exceptions were observed. For example, several NLRs located on chromosome 1 and chromosome 6 of the ‘124Pan’ genome were syntenic with chromosomes 2 and 1, respectively, in the ‘Lovell 2D’ v3.0 genome, suggesting instances of NLR translocation. Interchromosomal translocations of R-genes are often mediated by transposable element movement and can facilitate the introduction of novel integrated domains in NLRs that have an impact on function and resistance (Bailey et al., 2018; Krasileva, 2019).

**Figure 3.**
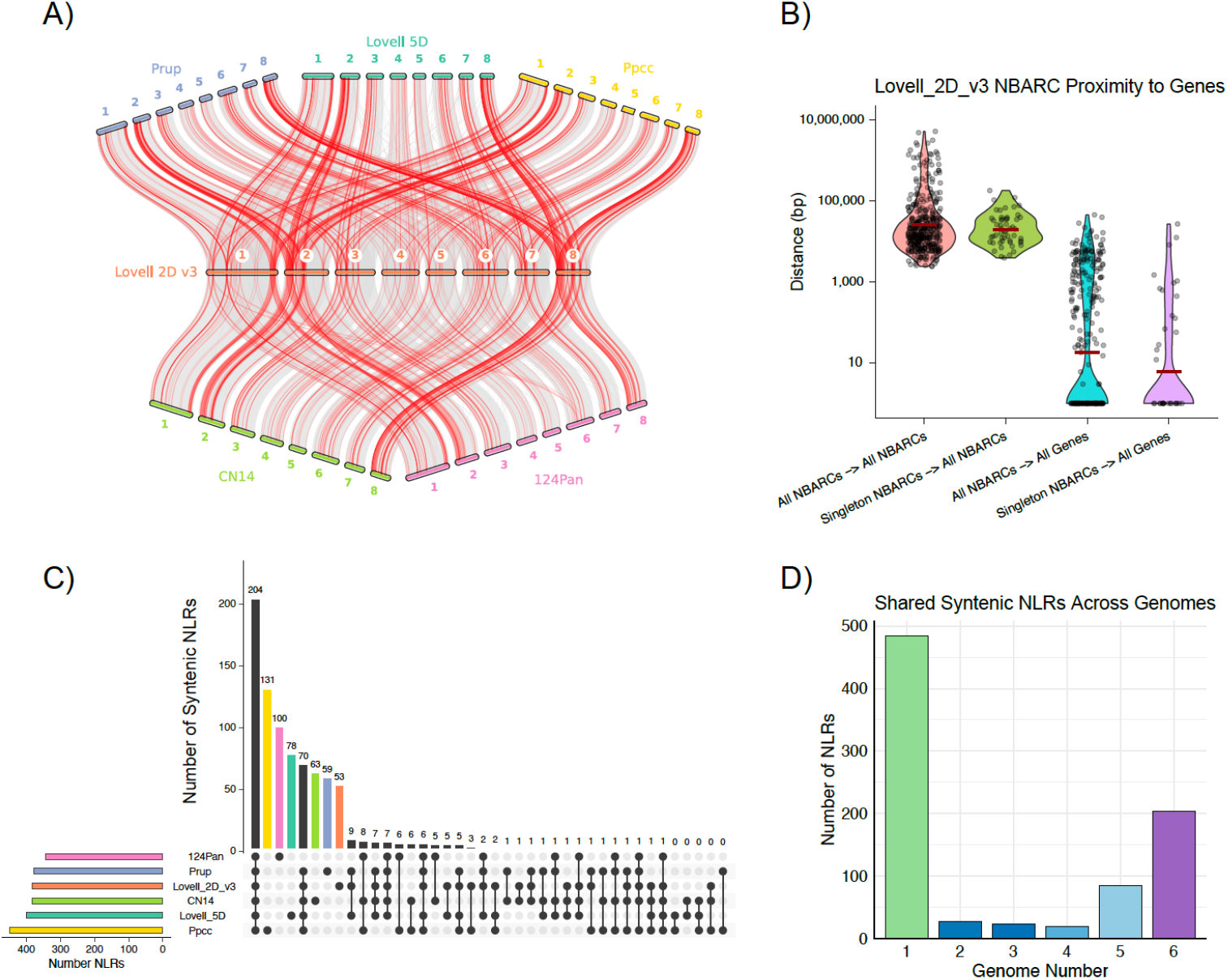
Pan-NLR analyses for the six genomes of *P. persica*. A) Macrosynteny (gray) of five P. persica genomes against the reference, Lovell 2D v3, along with syntenic orthologous NLR genes (red). B) Nearest NLR neighbors in Lovell 2D v3, with the distance to the nearest NBARC shown in red, the distance to the nearest NBARC for singletons (those NBARCs unique to Lovell 2D v3) in green, the distance of NBARCs to the nearest coding gene in blue, and the distance of singleton NBARCs to the nearest coding gene in purple. C) Upset plot of NBARCs in the pan-NLRome with those NBARCs unique to a single genome being colored according to that genome. D) Number of NBARCs found to be unique to a single genome (green), shared among 2 - 5 genomes (shades of blue), and those found to be core (found in all six genomes, purple).

We next examined the spatial distribution of NLR genes within the genome, focusing on proximity to other NLRs and annotated coding sequences. To do so, we calculated distances between each NBARC domain and its nearest neighboring NBARC, and also to the closest gene model from the structural annotation (Figure 3B). In general, NLR genes were found to be more closely positioned to annotated coding genes than to other NLRs. This trend was also observed among singleton NLRs (defined as NLRs present in only one genome and lacking syntenic orthologs in the remaining accessions) suggesting that these loci are not preferentially clustered and may not arise from recent tandem duplication events.

We compared the conservation of NBARC loci across the pan-NLRome and showed that the majority of NBARC loci are shared among two to five genomes, with a substantial subset (n = 204) classified as core, being conserved across all six accessions (Figure 3C, 3D). Singleton NLRs (those unique to a single genome) comprised 484 of the total NLRs identified, ranging from 53 in ‘Lovell 2D’ v3.0 to 131 in the ‘Chinese Cling’ genome (Figure 3D). These data indicate a large reservoir of genome-specific NLR diversity across *P. persica* accessions, underscoring the dynamic nature of resistance gene evolution even within a single species. Together, these insights provide a valuable framework for identifying candidate loci in future studies and for breeding programs aiming to harness the diversity of disease resistance genes for crop improvement.

**Table 4. *R*-gene classifications that were annotated using the FindPlantNLR pipeline in *P. persica* genomes**.

## Conclusion

In this study, we present improved chromosome-level genome assemblies for the doubled haploid peach genotypes ‘Lovell 2D’ and ‘Lovell 5D’, generated with Oxford Nanopore PromethION and PacBio HiFi. The resulting assemblies exhibit greatly increased contiguity and completeness, allowing more accurate representation of repetitive genomic regions compared to previously published peach genome resources. Additionally, the new annotations generated here provide improved gene models and detailed characterization of transposable elements and non-coding RNAs. Our comparative genomic analyses confirm high synteny across peach genomes but also revealed structural variation that may underlie phenotypic and agronomic differences. Further, in-depth characterization of the NLR gene repertoire across multiple peach accessions highlights the diversity within disease resistance loci, providing valuable insights for future breeding strategies aimed at developing resilient peach cultivars. Collectively, these improved genomic resources establish a foundation for future genetic, functional, and evolutionary studies, and will accelerate marker-assisted selection and breeding programs focused on sustainable peach production. Upon acceptance of the manuscript, the genome data will be integrated into GDR, allowing users to access marker, GWAS, and QTL data aligned to the genome sequences, along with additional functional annotations and syntenic regions across key Rosaceae genomes through search pages and graphical interfaces.

## Data Availability

Nanopore and HiFi sequence data will be deposited on to the NCBI SRA database once published. The genome assembly and gene annotations have been deposited on the GDR under accession number tfGDR1086 (Jung *et al*., 2019). Additionally, the genome and annotations for genes and NLRs have been deposited in a Zenodo Repository DOI: 10.5281/zenodo.15576776.

## Funding

Funding for the sequencing, assembly, and annotation were provided by appropriated funds award to CG and CD under USDA-ARS National Program 301 Plant Genetic Resource, Genomics and Genetic Improvement, project number 8080-21000-033-000D. Indexing, hosting, and further analysis of the genomic resources provide by GDR was made possible from funding awarded to SJ, PZ, and DM under SCRI-NIFA Award 2022-51181-38449.

## Conflict of Interest

The authors declare they have no conflicts of interest.

---

**Supplementary Figure 1.**
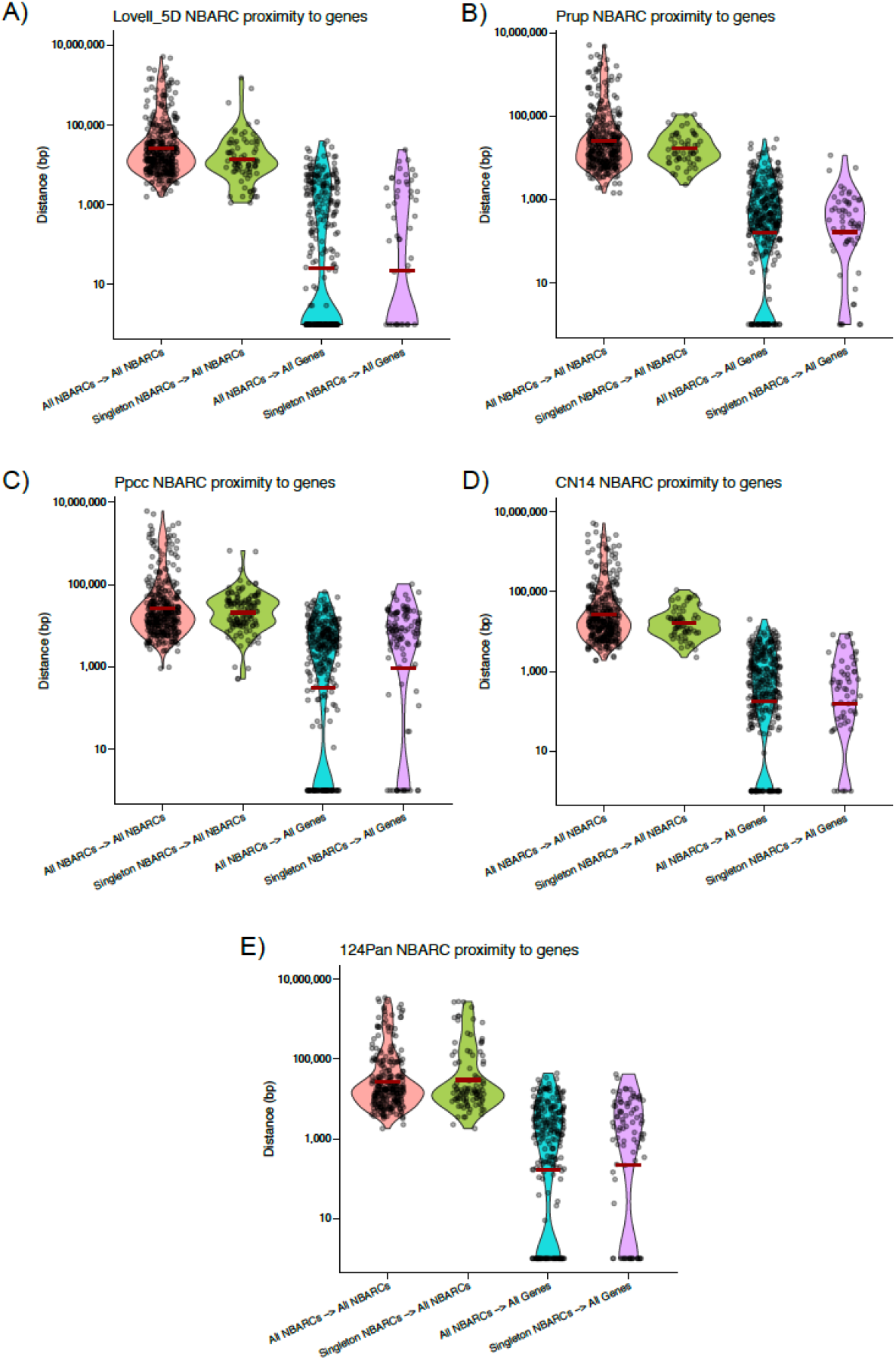
Nearest NLR neighbors in A) Lovell_5D, B) Prup, C) Ppcc, D) CN14, E) 124Pan, with the distance to the nearest NBARC shown in red, the distance to the nearest NBARC for singletons (those NBARCs unique to Lovell 2D v3) in green, the distance of NBARCs to the nearest coding gene in blue, and the distance of singleton NBARCs to the nearest coding gene in purple.

## Notes

### Competing Interest Statement

The authors have declared no competing interest.

